# *In vitro* antimicrobial and cytotoxic activity of *Lippia origanoides* essential oil against bacteria of potential health concern

**DOI:** 10.1101/2021.09.02.458771

**Authors:** Edwin Stiven Quiguanás-Guarín, Juan Pablo Bedoya Agudelo, Jhon Esteban López-Carvajal, Yuly Andrea Ramírez Tabares, Leonardo Padilla Sanabria, Jhon Carlos Castaño-Osorio

## Abstract

Due to the growing resistance they develop of bacteria to drugs, the search for alternatives in natural products is considered important such as Lippia origanoides essential oil. Here, the antibacterial activity of the oil and two of its major chemical components were tested against bacteria of potential health concern. The cytotoxicity of these compounds was evaluated in human erythrocytes and Vero cells. 51 compounds were identified in the LOEO, being terpinen-4-ol, γ-Terpinene, citronellal and thymol the main. LOEO and thymol showed antibacterial activity from 904 μg/mL and 200 μg/mL, respectively. γ-Terpinene did not show activity any concentration tested. LOEO showed hemolysis at concentration of 3000 μg/mL and thymol at 100 μg/mL. LOEO and thymol showed cytotoxicity in the evaluated cell lines at 250 μg/mL and 100 μg/mL, respectively. These compounds have a moderate cytotoxicity so it’s considered necessary to study alternatives to reduce the in vitro cytotoxicity of these compounds.

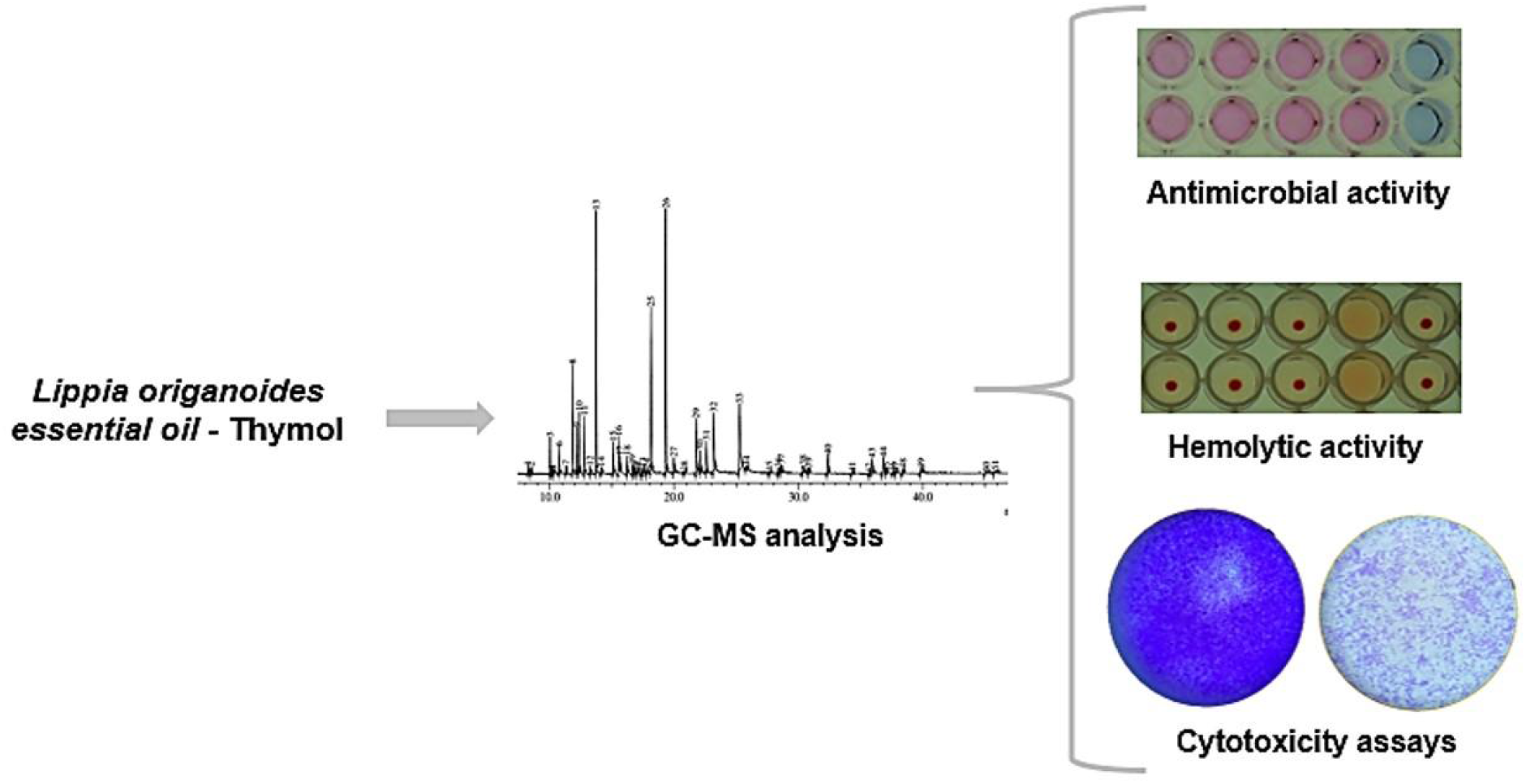

## 1. Introduction

Antimicrobial resistance (AMR) is a major growing threat to global public health, killing more than 700,000 people a year, but that number can increase to 10 million by 2050 (O’Neill 2014). The rapid adaptability of bacteria to new environmental conditions and the mobilization of resistance genes among them, imply that all antibiotics in clinical use may have bacteria that can resist them (Colzi et al. 2015).

Usually, infections caused by resistant bacteria are associated with increased morbidity and mortality making them challenging for therapeutic treatment (Walsh and Wencewicz 2014). Therefore, it’s necessary to implement therapeutic alternatives other than antibiotics (Alós 2015), such as natural products, because substances that are extracted from plants and other natural sources, can provide different compounds that are structurally diverse making resistance difficult (Chouhan et al. 2017). Several plants and herbs have been known for their antimicrobial properties and these properties are conferred by their essential oils (EO) (Azizkhani et al. 2012).

EOs are secondary metabolites obtained from aromatic plants, of a dense and volatile nature (Pandey et al. 2017). These oils are complex mixtures of volatile compounds biosynthesized by plants that include, terpenes, terpenoids, aromatic and aliphatic constituents, characterized by a low molecular weight (Pichersky et al. 2006). The mixture of their components causes an affectation to multiple chemical processes in bacteria, causing a considerable antibacterial effect (Bassolé and Juliani 2012) and therefore, can serve as a powerful tool to reduce antibacterial resistance (Stefanakis et al. 2013). The antimicrobial properties of EOs are well demonstrated against a wide spectrum of bacteria and the most of their effects appears to be related to phenolic compounds (Lv et al. 2011).

*Lippia origanoides* Kunth. species, belonging to the Verbenaceae family, produces an essential oil that has demonstrated several biological activities (Guimarães et al. 2015; Helal et al. 2019). *Lippia origanoides* essential oil (LOEO) has antioxidant, anti tumor and antimicrobial properties (Guimarães et al. 2015; Morão et al. 2016) and previous studies have confirmed this biological potential (Acosta et al. 2019). In addition, some of the oil’s major compounds have also shown similar biological activity (Sim et al. 2019; Sepahvand et al. 2021). Thus, in this study we evaluated the antibacterial activity of LOEO and two of the main chemical components against bacteria of sanitary importance, also, the cytotoxicity of the compounds was evaluated in three cell lines.

## 2. Results and Discussion

### 2.1. Chemical analysis of LOEO by GC-MS

The chemical composition of the LOEO was elucidated by gas chromatography coupled to mass spectrometry (GC-MS) and it was possible to determine a total of 51 different chemical compounds, of which monoterpenes such as Terpinen-4-ol (17.84%), γ-Terpinene (14.72%), citronellal (10.08%) and Thymol (7.28%) were found in a higher proportion (Table 1). For thymol, similar results have been evidenced in LOEO, where it has been reported as a main component (Barreto et al. 2014; Cáceres et al. 2020).

**Table 1.** Chemical constituents of *Lippia origanoides* essential oil.

| Compound | Area % | Compound | Area % |
| --- | --- | --- | --- |
| 1. .alpha.-Phellandrene | 0,28 | 27. L-.alpha.-Terpineol | 1,03 |
| 2. .alpha.-Pinene | 0,27 | 28. 2-Cyclohexen-1-ol, 3-methyl-6-(1-methylethyl) | 0,13 |
| 3. Bicyclo[3.1.0]hexane, 4-methylene-1-(1-methylethyl) | 1,78 | 29. Citronellol | 3,88 |
| 4. .beta.-Pinene | 0,14 | 30. Benzene, 2-methoxy-4-methyl-1-(1-methylethyl) | 1,46 |
| 5. 1-Octen-3-ol | 0,24 | 31. Benzene, 2-methoxy-4-methyl-1-(1-methylethyl) | 2,27 |
| 6. .beta.-Myrcene | 1,39 | 32. Geraniol | 5,28 |
| 7. .alpha.-Phellandrene | 0,37 | 33. Thymol | 7,28 |
| 8. Cyclohexene, 1-methyl-4-(1-methylethylidene) | 5,83 | 34. Phenol, 2-methyl-5-(1-methylethyl) | 0,27 |
| 9. p-Cymene | 2,41 | 35. .gamma.-Elemene | 0,16 |
| 10. D-Limonene | 3,44 | 36. 3-Cyclohexene-1-methanol, .alpha.,.alpha.,4-trimet | 0,46 |
| 11. trans-.beta.-Ocimene | 3,05 | 37. 2,6-Octadiene, 2,6-dimethyl- | 0,71 |
| 12. 1,3,6-Octatriene, 3,7-dimethyl-, (Z) | 0,32 | 38. 2,6-Octadien-1-ol, 3,7-dimethyl-, acetate, (Z) | 0,89 |
| 13. .gamma.-Terpinene | 14,72 | 39. Cyclohexane, 1-ethenyl-1-methyl-2,4-bis(1-methylet) | 0,45 |
| 14. p-Menth-8-en-1-ol, stereoisomer | 0,37 | 40. Bicyclo[7.2.0]undec-4-ene, 4,11,11-trimethyl-8-met) | 1,7 |
| 15. Cyclohexene, 1-methyl-4-(1-methylethylidene) | 1,89 | 41. Humulene | 0,09 |
| 16. p-Menth-8-en-1-ol, stereoisomer | 1,84 | 42. Naphthalene, 1,2,4a,5,6,8a-hexahydro-4,7-dimethyl | 0,04 |
| 17. 1,6-Octadien-3-ol, 3,7-dimethyl | 0,53 | 43. Naphthalene, 1,2,4a,5,6,8a-hexahydro-4,7-dimethyl | 1,41 |
| 18. 1-Octen-3-yl-acetate | 0,99 | 44. .gamma.-Elemene | 1,57 |
| 19. 2-Cyclohexen-1-ol, 1-methyl-4-(1-methylethyl) | 0,57 | 45. .alpha.-Muurolene | 0,06 |
| 20. 3-Tetradecanol acetate | 0,07 | 46. .beta.-Bisabolene | 0,06 |
| 21. 2,4,6-Octatriene, 2,6-dimethyl | 0,13 | 47. .gamma.-Muurolene | 0,1 |
| 22. Bicyclo[3.2.1]oct-2-ene, 3-methyl-4-methylene | 0,02 | 48. Naphthalene, 1,2,3,5,6,8a-hexahydro-4,7-dimethyl-1 | 0,38 |
| 23. 2-Cyclohexen-1-ol, 1-methyl-4-(1-methylethyl) | 0,41 | 49. Cyclohexanemethanol, 4-ethenyl-.alpha.,.alpha.,4-t | 0,54 |
| 24. Isopulegol | 0,36 | 50. 1-Naphthalenol, 1,2,3,4,4a,7,8,8a-octahydro-1,6-di | 0,18 |
| 25. Citronellal | 10,08 | 51. 1H-Cycloprop[e]azulen-4-ol, decahydro-1,1,4,7-tetr | 0,26 |
| 26. Terpinen-4-ol | 17,84 | **Total** | **100** |

### 2.2. Antimicrobial activity of Essential oil and their main components

DMSO was used to solubilize LOEO, thymol and γ-Terpinene. But first of all, the minimum inhibitory concentration (MIC) of DMSO was tested for each bacterium so it wouldn’t affect the growth of the microorganisms (table 2), reaching a consensus concentration of 5% DMSO as a vehicule.

**Table 2.** Minimum Inhibitory Concentration (MIC) in μg/mL of the LOEO and their main components.

| Bacteria | DMSO % | LOEO<br>(µg/mL) | Thymol<br>(µg/mL) | γ-Terpinene<br>(µg/mL) |
| --- | --- | --- | --- | --- |
| *Acinetobacter* spp ATCC 49139 | 9 | 1.808 | 300 | - |
| *Escherichia coli* ATCC 25922 | 8 | 1.808 | 300 | - |
| *Klebsiella pneumoniae* ATCC 13882 | 10 | 1.808 | 200 | - |
| *Staphylococcus aureus* ATCC 25923 | >10 | 5.424 | 400 | - |
| *Shiga toxin-producing Escherichia coli* (103) | 8 | 904 | 250 | - |
| *Shiga toxin-producing Escherichia coli* (106) | 8 | 1.808 | 300 | - |
| *Shiga toxin-producing Escherichia coli* (N3) | 8 | 1.808 | 300 | - |

Antimicrobial activity of LOEO, thymol and γ-Terpinene was evaluated against both ATCC reference strains and bovine isolates. The LOEO showed antimicrobial activity against *Acinetobacter* spp 49139, *Escherichia coli* 25922, *Klebsiella pneumoniae* 13882 ATCC with MICs around 1,808 μg/mL and 904 to 1,808 μg/mL against 103, 106 and N3 STEC bovine isolate, respectively. However, LOEO showed antimicrobial activity at higher concentration against *Staphylococcus aureus* 25923 (MIC 5,424 μg/mL). This effect has been evidenced in other studies; for example, Perera et al. (2016) report an antibacterial effect of LOEO at 625 μg/mL against *Escherichia coli* ATCC 11229; however, they did not show activity against *Staphylococcus aureus* (MRSA BMB 9393).

For Shiga toxin-producing *Escherichia coli* (STEC) strains, we report MIC of the LOEO of 904 and 1808 μg/mL (0.1 and 0.2%); in contrast, Lizcano et al. (2020), reported antibacterial activity of LOEO at a concentration of 1.8% against these same bacteria. Besides, Caceres et al. (2020), reported antibacterial activity of two chemotypes of the *Lippia origanoides* essential oil at concentrations of 370 and 750 μg/mL against *E. coli* O33 and O157:H7. Essential oils such as LOEO can affect the cytoplasm and membranes of bacteria, therefore, the mechanisms of action of this type of oil include cell wall degradation and damage to membrane proteins, reducing synthesis of ATP (da Cunha et al. 2018).

Thymol showed antimicrobial activity in the bacteria evaluated at concentrations ranging between 250 and 400 μg/mL. Similar results have been reported in the characterization of other essential oils. For example, Flores et al. (2014), identified thymol as one of the main compounds of the essential oil of *Thymus vulgaris* and showed an antimicrobial effect at concentrations of 2.540 μg/mL against *Escherichia coli* strains. Overall, it is assumed that the biological potential of essential oils is attributed to the main chemical compounds, in this case, terpenoids such as thymol, have antibacterial activity mediated by the functional group that acts on the outer membrane of bacteria, altering thus the permeability and/or fluidity of the membranes and affecting their proteins and periplasmic enzymes (da Cunha et al. 2018), which causes cell death.

Nonetheless, γ-Terpinene did not show antimicrobial activity at any concentration tested (100 to 900 μg/mL) (Table 2). Dorman and Deans (2000), cite that although this compound has been found in considerable amounts in the essential oil of *Thymus vulgaris*, it has not been associated with antibacterial properties. However, the possibility that less abundant molecules in these types of oils are more effective than the main compounds should not be ruled out (Yap et al. 2014).

### 2.3. Compounds Minimum Cytotoxic Concentration (CC50), Inhibitory Concentration (IC50), and Therapeutic Index (TI)

Compounds toxicity in mammalian cells was evaluated in human erythrocytes and Vero cell line (Figure 1). For LOEO, concentrations ranging from 125 to 3000 μg/mL were tested. LOEO showed cytotoxic activity to concentrations from 250 to 3000 μg/mL, while thymol showed toxicity to concentrations from 100 to 400 μg/mL. Defining the possible cytotoxic effects of essential oils is an important issue considering their possible use as a therapeutic alternative (Mohmod et al. 2015), therefore, when evaluating the cytotoxicity of the compound, a CC50 = 230.2 μg/mL and a CC50 = 63.56 μg/mL were evidenced for LOEO and thymol respectively (Table 3).

**Figure 1.**
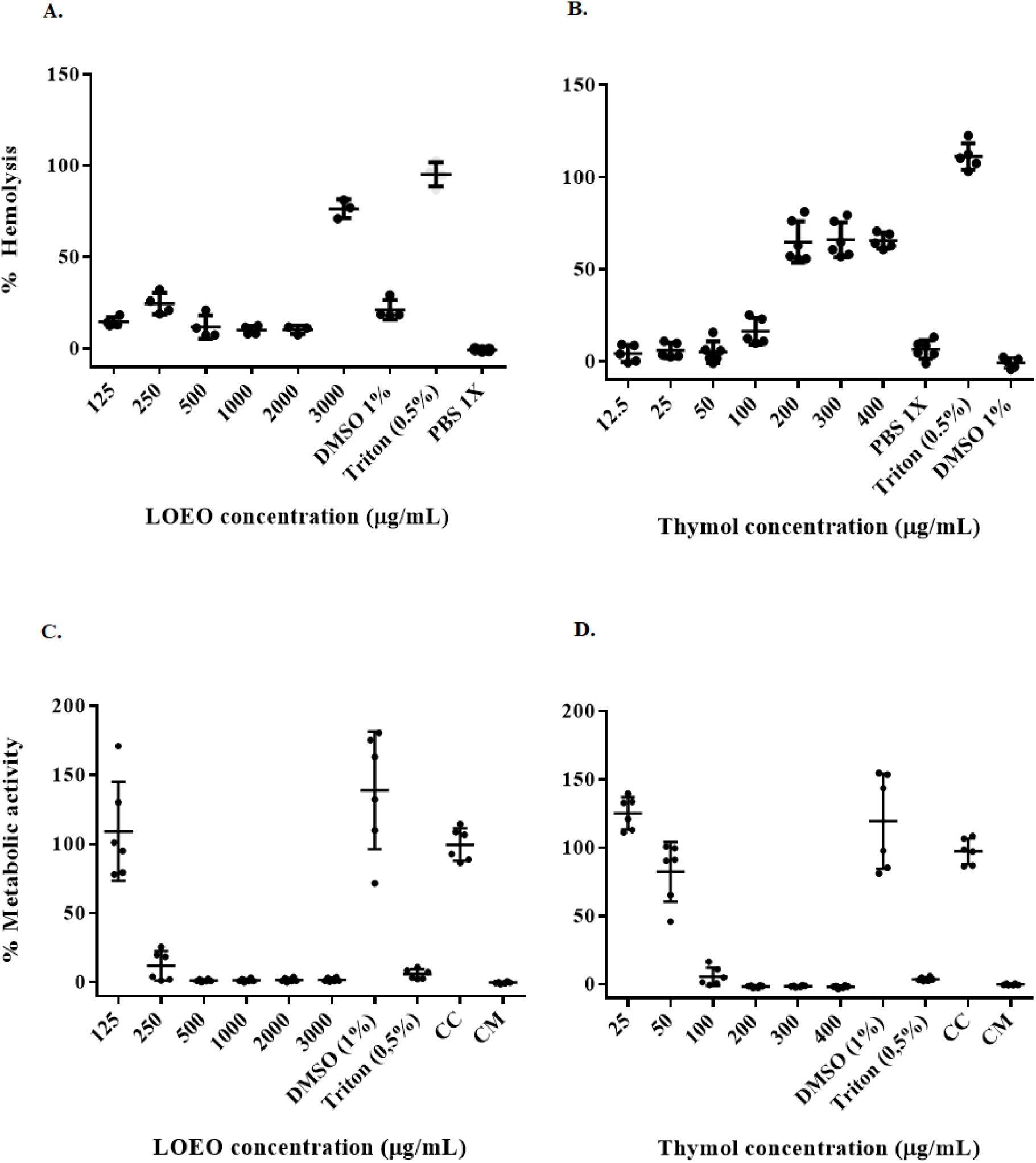
Compounds toxicity against line cells. LOEO (A) and thymol (B) toxicity against human erythrocytes. LOEO (C) and thymol (D) toxicity against Vero cells. Data shown as mean with standard deviation (n = 6). CC = Cells control. CM = control medium DMEM.

**Table 3.**
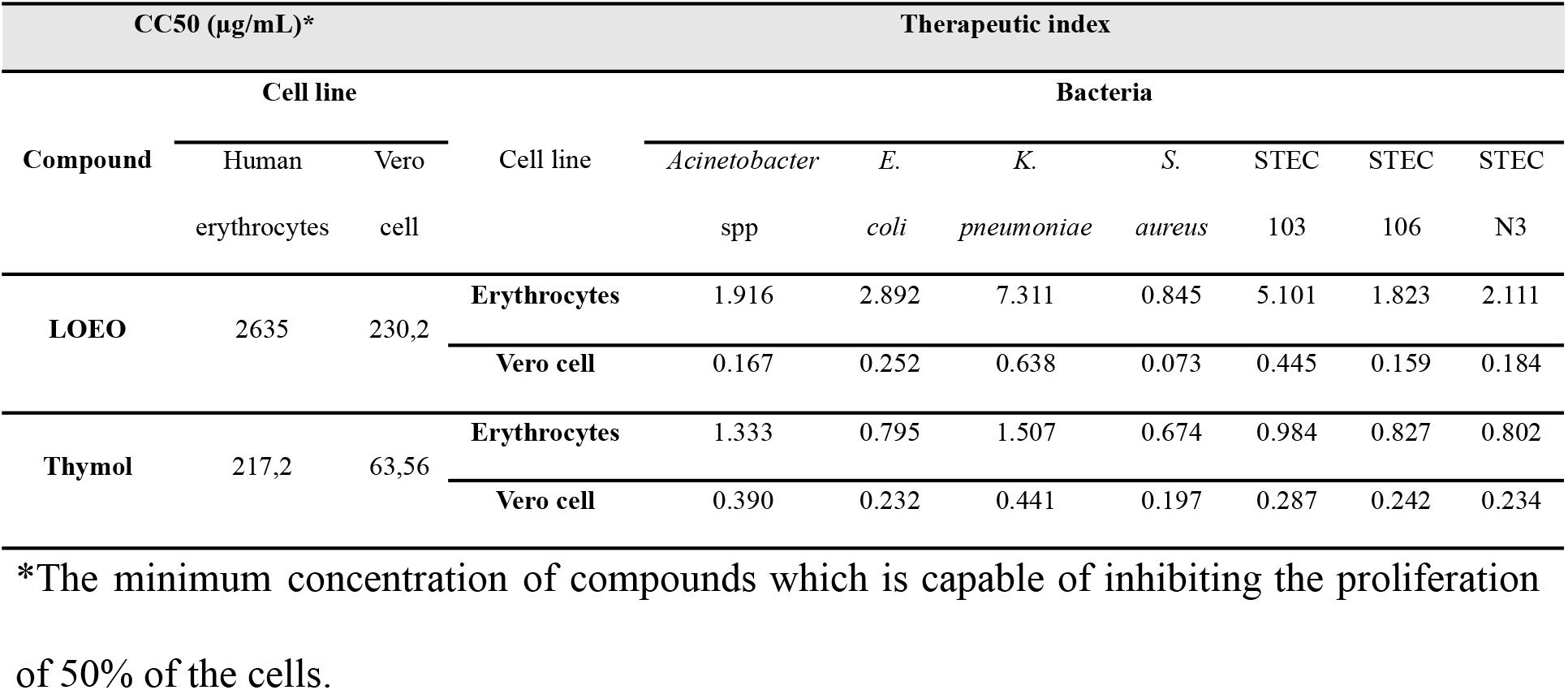
Minimum Cytotoxic Concentration, Inhibitory Concentration, and Therapeutic Index of LOEO and Thymol.

| CC50 (µg/mL)* |  |  | Therapeutic index |  |  |  |  |  |  |  |
| --- | --- | --- | --- | --- | --- | --- | --- | --- | --- | --- |
| Compound | Cell line |  | Cell line | Bacteria |  |  |  |  |  |  |
|  | Human erythrocytes | Vero cell |  | *Acinetobacter* spp | *E. coli* | *K. pneumoniae* | *S. aureus* | STEC 103 | STEC 106 | STEC N3 |
| LOEO | 2635 | 230,2 | Erythrocytes | 1.916 | 2.892 | 7.311 | 0.845 | 5.101 | 1.823 | 2.111 |
|  |  |  | Vero cell | 0.167 | 0.252 | 0.638 | 0.073 | 0.445 | 0.159 | 0.184 |
| Thymol | 217,2 | 63,56 | Erythrocytes | 1.333 | 0.795 | 1.507 | 0.674 | 0.984 | 0.827 | 0.802 |
|  |  |  | Vero cell | 0.390 | 0.232 | 0.441 | 0.197 | 0.287 | 0.242 | 0.234 |

\*The minimum concentration of compounds which is capable of inhibiting the proliferation of 50% of the cells.

Ríos et al. (2008), mentioned that essential oils with CC50 ≤ 100 μg/mL are considered to have powerful cytotoxic activity; CC50 values between 100 and 500 μg/mL are considered moderately toxic; Values between 500 and 1000 μg/mL are considered weak cytotoxic and values above 1000 μg/mL are classified as non-toxic to mammalian cells. Thus, based on these criteria, LOEO presented moderate cytotoxicity against Vero cells, while thymol presented powerful cytotoxic activity.

The CC50 for cell lines and IC50 for each bacteria were evaluated using the GraphPad Prism 7 statistical software package. The data were used to determine the TI (TI = CC50 / IC50). The interpretation was done as follows, TI > 1 = low cytotoxicity and TI < 1 = High cytotoxicity (Muller and Milton 2012).

The results here reported agree with that indicated by Borges et al. (2012), who report moderate toxicity of the *Lippia sidoides* and *Lippia origanoides* essential oil in mammalian cells, with CC50 = 192.7 μg/mL and CC50 = 175.7 μg/mL respectively. In addition, Cáceres et al., 2020, reported CC50 = 480 μg/mL and CC50 = 830 μg/mL respectively for two chemotypes (thymol-carvacrol) of *Lippia origanoides* essential oil. Finally, assessing the hemolytic activity contributes to the cytotoxicity of some compounds in mammalian cells (Zohra and Fawzia 2014). Therefore, in this study, LOEO was found to be less hemolytic (CC50 = 2635 μg/mL), compared to thymol, with CC50 = 217.2 μg/mL against human erythrocytes (Table 3).

## 3. Materials and methods

### 3.1. Essential oil and components

The LOEO was acquired from Natuaroma company. The density of the oil was determined by weighing 1 mL in triplicate in sterile 1.5 mL tubes on a Shimadzu AUY120 analytical balance. Thymol (98.5%, Sigma Aldrich T0501-500g) and γ-Terpinene (97 %, Sigma Aldrich 223190) were diluted in 5% DMSO to obtain a stock solution of 10 mg/mL for each compound.

### 3.2. Bacterial Strains

The following bacteria were used in this study: *Escherichia coli* (ATCC 25922), *Klebsiella pneumoniae* (ATCC 13882), Acinetobacter spp. (ATCC 49139), *Pseudomonas aeruginosa* (ATCC 9027) and *Staphylococcus aureus* (ATCC 25923). In addition, three isolates of STEC from livestock feces were included (Quiguanás-Guarín et al. 2021). Initially, all microorganisms were seeded on Müller Hinton agar medium (MHA) (Scharlau 01-136-500, Barcelona, Spain) and incubated at 37 °C for 24 hours, and then, one colony of each microorganism was subcultured in Müller Hinton broth medium (MHB) (Scharlau 02-136-500, Barcelona, Spain) at 37 °C for 16 hours.

### 3.3. Gas chromatography-mass spectrometry (GC-MS) analysis of LOEO

The analysis of LOEO was carried out by GC-MS using Shimadzu equipment (Injector: AOC-20i; Sampler: AOC-20s; GC: GC-2010 Plus and GCMS-QP2010 Ultra), equipped with a flame ionization detector (FID). The sample was dissolved in chloroform (SupraSolv Merck 1-024321000) at a concentration of 1 mg/mL. 1 μL of sample was injected in split injection mode (1:10 at 250 °C) with a flow rate of 1mL/min in a Rxi-5ms column (RESTEK) (Crossbond 5% dipheniyl / 95% dimethyl polysiloxane) with dimensions 30m x 0.25mm ID, 0.25um df, chromatographic grade helium (99.9999% purity) was used as carrier gas.

The following temperature gradient was used: 1 min at 50 °C, increasing 3 °C/min to 110 °C for 1 minute; it increased 2 °C/min to 200 °C for 1 minute and finally increased 10 ° C/min to 250 °C for five minutes. The detector in electronic impact mode (70eV) started after 3 min of injection, scanning masses between 35 and 500 m/z every 0.30 seconds.

### 3.4. Antimicrobial activity assay

Antimicrobial activity of LOEO, thymol and γ-Terpinene was tested by the broth microdilution assay as described by Wiegand et al. (2008) with some modifications.

The bacteria were grown overnight in MHA and then adjusted to an absorbance of 0.4 (A570 nm) (3-5 × 10^8^ CFU/mL). The bacteria inoculum were adjusted to a final dilution of 1:1000 in MHB (3-5 × 10^5^ CFU/mL); 90 μL of bacteria, and 10 μL of each compound were mixed to a final concentration between 452 - 9,040 μg/mL for LOEO and 100 - 900 μg/mL for thymol and γ-Terpinene. The solution was added in 96-well microplates and incubated at 37 °C for 16 h and resazurin (Redox indicator, Acros Organics 418900050, Geel, Belgium) was added to a final concentration of 44 μM; the plate was incubated for an additional 1 h and then the fluorescence values at 565/600 nm excitation/emission were measured in plate reader (Synergy HTX, Biotek). The average value of the blanks was subtracted from each sample and the growth percentage was calculated relative to a growth control. Each concentration was measured in triplicate and the MIC was defined as the lowest concentration (in μg/mL) that inhibited 50% growth in bacteria after 20 hours of incubation

### 3.5. Human erythrocyte hemolytic activity

Three milliliters of human heparinized or with EDTA blood were centrifuged at 800 g for 10 min at room temperature. The erythrocytes were washed three times with a 1X PBS stock solution (130 mM NaCl, 3 mM KCl, 8 mM Na_2_HPO_4_, 1.5 mM K2HPO4, pH 7.4). The supernatant was discarded and 1X PBS was added in a proportion that will be equal to the initial volume of blood. An erythrocyte dilution at 1:250 was prepared from the erythrocyte stock solution and incubated at 37 °C for 15 min (work solution). In 96-well polypropylene microplates (Greiner bio-one 650201, USA), were added 90 μL of erythrocyte solution and 10 μL of LOEO solutions to final concentrations ranging from 125 to 3000 μg/mL and between 12.5 and 400 μg/mL for Thymol, to a final volume of 100μL per well. Untreated cells and cells treated with Triton X-100 0.05 % were included as controls.The erythrocytes were then incubated for 18 hours at 37 °C and the plate was centrifuged at 800 g for 5 min a room temperature and without brake in the centrifuge Thermo Scientific Heraeus Megafuge 11. Supernatants (80 μL per well) were carefully taken and transferred to another polystyrene plate to measure hemoglobin release by its absorbance at 540 nm in an spectrophotometer (EPOCH Biotek). The absorbance of the hemoglobin from the erythrocytes incubated with Triton-X100 was taken as 100% hemolysis (Toro et al. 2017).

### 3.6. Cytotoxicity assays of compounds

The cytotoxic activity of the compounds was evaluated in Vero cells (ATCC-CLL 81) using a modified microdilution assay (O’Brien et al. 2000). Vero cells were cultured in Dulbecco’s modified Eagle medium (DMEM) - (D5648 Sigma) supplemented with 5% (v/v) Fetal Bovine Serum (FBS) (CVFSVF00-01 Eurobio), 1X antibiotic antimycotic (sigma A5955), and 2 mM L-glutamine (GLL01-100ML caisson), and maintained at 37 °C 5% CO_2_ atmosphere. 15,000 Vero cells were seeded in 96-microwell plates and incubated with the compounds at final concentrations ranging from 125 to 3000 μg/mL or 12.5 to 400 μg/mL of essential oil or thymol, respectively. After 18-24 h of incubation, resazurin was added to a final concentration of 44 μM and incubated for 4 hours.

The metabolic activity was measured by the converted resorufin fluorescence at 565/10 nm (ex.) and 600/40 nm (em.) in a Synergy HTX (biotek) fluorometer. Untreated cells (CC) and cells with 0.5 % Triton X-100 (CT) were included as controls. For the analysis of results, fluorescence intensity was used to calculate the metabolic activity (X*100/CC). Minimal toxic concentrations were defined as the minimal compounds concentration that showed a statistically significant effect compared with the cell control (Téllez and Castaño-Osorio 2014).

### 3.7. Inhibitory concentration (IC50), and therapeutic index (TI)

The toxic and antimicrobial activity inhibitory concentration (IC50), was calculated using sigmoidal dose-response curves from the active compounds with the GraphPad Prism 7 program. The TI was calculated with the ratio of the toxic IC50 and the antimicrobial activity IC50 for each compound using experimental data from cytotoxicity in human erythrocytes and VERO (ATCC^®^ CCL-81) cells concerning their IC50 in the antimicrobial activity against bacteria.

### 3.8. Statistical analysis

The experimental data were analyzed using One-way ANOVA, Dunnett’s multiple comparisons test in the statistical software package of GraphPad Prism 7 program. P values less than 0.05 were considered statistically significant. All data were presented as mean ± standard deviation and were the results of at least two independent experiments with triplicate assays.

## 4. Conclusion

Essential oils are considered an alternative to be used as natural antimicrobials against different bacterial species. However, these compounds have a moderate to high cytotoxicity, so it’s considered necessary to study alternatives to reduce the *in vitro* cytotoxicity of these compounds. Overall, these results show a promising path in the development of therapeutic strategies for antimicrobial resistance problems.

## Confidentiality of the data

The authors declare that they have followed the protocols of your work center on the publication of patient data.

## References

Acosta JM, Arango O, Álvarez DE, Hurtado, AM. 2019. Actividad Biocida del Aceite Esencial de *Lippia origanoides* H.B.K sobre Phytophthora infestans (Mont.) de Bary [Biocidal Activity of the Essential Oil of Lippia origanoides H.B.K on Phytophthora infestans (Mont.) of Bary]. Inf Tecnol. 30(6): 45–54 Spanish.

Alós J-I. 2015. Resistencia bacteriana a los antibióticos: una crisis global [Bacterial resistance to antibiotics: a global crisis]. Enfer Infec y Micro Clínica. 33(10):692–699 Spanish.

Azizkhani M, Basti AA, Misaghi A, Tooryan F. 2012. Effects of *Zataria multiflora* Boiss., *Rosmarinus officinalis* L. and *Mentha longifolia* L. essential oils on growth and gene expression of enterotoxins C and E in *Staphylococcus aureus* ATCC 29213. J. Food Saf. 32:508–516.

Barreto HM, de Lima IS. Nunes KM, Osório LA, Mourão RA, dos Santos BH, Coutinho HD, Lira de Abreu AP, de Medeiros MG, Lopes MG, et al. 2014. Effect of *Lippia origanoides* H.B.K. essential oil in the resistance to aminoglycosides in methicillin resistant *Staphylococcus aureus*. Eur J of Integr Med. 6(5):560–564

Bassolé IH, Juliani HR. 2012. Essential oils in combination and their antimicrobial properties. Molecules 17: 3989–4006

Borges AR, Aires JR, Maciel TM, Freire M, Graças AM, Dantas JA, Queiroz RC. 2012. Trypanocidal and cytotoxic activities of essential oils from medicinal plants of Northeast of Brazil. Exp Parasitol. 132(2):123–128

Caceres M, Hidalgo W, Stashenko E, Torres R, Ortiz C. 2020. Essential oils of aromatic plants with antibacterial, anti-biofilm and anti-quorum sensing activities against pathogenic bacteria. Antibiotics. 9:147.

Chouhan S, Sharma K, Guleria S. 2017. Antimicrobial activity of some essential oils—present status and future perspectives. Medicines. 4(3):58.

Colzi I, Troyan AN, Perito B, Casalone E, Romoli R, Pieraccini G, Škalko-Basnet N, Adessi A, Rossi F, Gonnelli C, et al. 2015. Antibiotic delivery by liposomes from prokaryotic microorganisms: Similia cum similis works better. Eur J of Phar and Bioph. 94(39): 411–418

da Cunha JA, Heinzmann BM, Baldisserotto B. 2018. The effects of essential oils and their major compounds on fish bacterial pathogens-a review. J of Appl Microbiol. 125(2):328–344

Dorman HJ, Deans SG. 2000. Antimicrobial agents from plants: antibacterial activity of plant volatile oils. J. Appl. Microbiol. 88:308–316

Guimarães LG, da Silva ML, Reis PC, Costa MT, Alves LL. 2015. General characteristics, phytochemistry and pharmacognosy of *Lippia sidoides*. Nat. Prod. Commun. 10, 1861–1867.

Helal IM, El-Bessoumy A, Al-Bataineh E, Joseph MR, Rajagopalan P, Chandramoorthy CH, Hadj SB. 2019. antimicrobial efficiency of essential oils from traditional medicinal plants of asir region, Saudi Arabia, over drug resistant isolates. BioMed Reser Inter. 29: 8928306.

Lizcano A, Villamizar R, Herrera F, Santos J. 2020. Inactivation of Shiga Toxin-producing Escherichia coli in meat under refrigerated conditions using the essential oil of *Lippia origanoides*. Bistua: Rev de la Fac de Cienc Básicas. 18:27–33 Spanish.

Lv F, Liang H, Yuan Q, Li, C. 2011. In vitro antimicrobial effects and mechanism of action of selected plant essential oil combinations against four food-related microorganisms. Food Res. Int. 44: 3057–3064.

Mohmod AL, Krishnasamy G, Adenan MI. 2015. Malaysian plants with potential in vitro trypanocidal activity. Ann. Phytomed. 4:6–16.

Morão RP, de Almeida AC, Martins ER, Bicalho JP, de Oliveira FD. 2016. Constituintes químicos e princípios farmacológicos do óleo essencial de alecrim pimenta (*Lippia origanoides*) [Constituents’ chemicals and principles pharmacological oil rosemary essential pepper (*Lippia origanoides*)]. Mt Claros. 18:74–81 Portuguese.

Muller PY, Milton MN. 2012. The determination and interpretation of the therapeutic index in drug development. Nat rev drug discov. 11(10):751–761

O’Brien J, Wilson I, Orton T, Pognan F. 2000. Investigation of the Alamar Blue (resazurin) fluorescent dye for the assessment of mammalian cell cytotoxicity: Resazurin as a cytotoxicity assay. Eur. J. Biochem. 267:5421–5426.

O’Neill, J. (2014). Antimicrobial Resistance: Tackling a crisis for the health and wealth of nations. London: Review on Antimicrobial Resistance. 2014. Available from: https://amr-review.org/sites/default/files/AMR%20Review%20Paper%20-%20Tackling%20a%20crisis%20for%20the%20health%20and%20wealth%20of%20nations_1.pdf

Pandey AK, Kuma P, Singh P, Tripathi NN, Bajpai VK. 2017. Essential oils: sources of antimicrobials and food preservatives. Front. Microbiol. 7:2161.

Perera WH, Bizzo HR, Gama PE, Alviano CS, Salimena FR, Alviano DS, Leitão SG. 2016. Essential oil constituents from high altitude Brazilian species with antimicrobial activity: *Baccharis parvidentata* Malag., *Hyptis monticola* Mart. ex Benth. and *Lippia origanoides* Kunth. J of Essent Oil Res. 29(2):109–116

Pichersky E, Noel JP, Dudareva N. 2006. Biosynthesis of plant volatiles: Nature’s diversity and ingenuity. Science. 311:808–811

Quiguanás-Guarín E, Granobles-Velandia C, Arango-Gil B, Giraldo-Rubio V, Cataño-Osorio J. 2021. Aislamiento de *Escherichia coli* productora de toxina Shiga (STEC) en heces de ganado y detección de factores de virulencia asociados con su patogénesis. Infectio. 25(1): 33–38

Ríos YK, Otero AC, Muñoz DL, Echeverry M, Robledo SM, Yepes MA. 2008. Actividad citotóxica y leishmanicida in vitro del aceite esencial de manzanilla (*Matricaria chamomilla*) [In vitro cytotoxic and leishmanicidal activity of chamomile essential oil (*Matricaria chamomilla*)]. Rev Col Cienc Quím Farm. 37:200–211.

Santurio FD, Jesus FP, Zanette RA, Schlemmer KB, Fratton A, Fries LL. 2014. Antimicrobial Activity of the Essential Oil of Thyme and of Thymol against Escherichia coli Strains. Acta Sci Vet. 42:1234.

Sepahvand S, Amiri S, Radi M, Akhavan H-R. 2021. Antimicrobial activity of thymol and thymol-nanoemulsion against three food-borne pathogens inoculated in a sausage model. Food Bioprocess Technol. [accessed 2021 Aug 16]:[6 p.]. https://doi.org/10.1007/s11947-021-02689-w

Sim JX, Khazandi M, Chan WY, Trott DJ, Deo TP. 2019. Antimicrobial activity of thyme oil, oregano oil, thymol and carvacrol against sensitive and resistant microbial isolates from dogs with otitis externa. Vet Dermatol. 30:524–e159.

Stefanakis M, Touloupakis E, Anastasopoulos E, Ghanotakis D, Katerinopoulos H, Makridis P. 2013. Antibacterial activity of essential oils from plants of the genus *Origanum*. Food Cont. 34(2):539–546

Téllez GA, Castaño-Osorio JC. 2014. Expression and purification of an active cecropin-like recombinant protein against multidrug resistance *Escherichia coli*. Protein Expr and Purif. 100: 48–53.

Toro LJ, Téllez GA, Henao CD, Rivera JD, Bedoya JP & Castaño-Osorio, JC. 2017. Identification and characterization of novel cecropins from the *Oxysternon conspicillatum* neotropic dung beetle. PLoS ONE. 12(11):1–17.

Walsh CT, Wencewicz TA. 2014. Prospects for new antibiotics: a molecule-centered perspective. The J of Antib. 67:7–22

Wiegand I, Hilpert K, Hancock RE. Agar and broth dilution methods to determine the minimal inhibitory concentration (MIC) of antimicrobial substances. Nat protoc. 3:163–75.

Yap PS, Yiap BC, Ping HC, Lim HE. 2014. Essential oils, a new horizon in combating bacterial antibiotic resistance. Open Microbiol. J. 8:6–14.

Zohra M, Fawzia A. 2014. Hemolytic activity of different herbal extracts used in Algeria. Int. J. Pharm. Sci. Res. 5:8494–8495.

